# DeepSelectNet: Deep Neural Network Based Selective Sequencing for Oxford Nanopore Sequencing

**DOI:** 10.1101/2022.10.24.513498

**Authors:** Anjana Senanayake, Hasindu Gamaarachchi, Damayanthi Herath, Roshan Ragel

## Abstract

**Background:** Nanopore sequencing allows selective sequencing, the ability to programmatically reject unwanted reads in a sample. Selective sequencing has many present and future applications in genomics research and the classification of species from a pool of species is an example. Existing methods for selective sequencing for species classification are still immature and the accuracy highly varies depending on the datasets. For the five datasets we tested, the accuracy of existing methods varied in the range of ~77%-97% (average accuracy <89%). Here we present DeepSelectNet, an accurate deep-learning-based method that can directly classify nanopore current signals belonging to a particular species. DeepSelectNet utilizes novel data preprocessing techniques and improved neural network architecture for regularization.

**Results:** For the five datasets tested, DeepSelectNet’s accuracy varied between ~91%-99% (average accuracy ~95%). At its best performance, DeepSelectNet achieved a nearly 12% accuracy increase compared to its deep learning-based predecessor SquiggleNet. Furthermore, precision and recall evaluated for DeepSelectNet on average were always >89% (average ~95%). In terms of execution performance, DeepSelectNet outperformed SquiggleNet by ~13% on average. Thus, DeepSelectNet is a practically viable method to improve the effectiveness of selective sequencing.

**Conclusions:** Compared to base alignment and deep learning predecessors, DeepSelectNet can significantly improve the accuracy to enable real-time species classification using selective sequencing. The source code of DeepSelectNet is available at https://github.com/AnjanaSenanayake/DeepSelectNet.

## Background

### Problem

Oxford Nanopore Technologies (ONT) introduced *MinION*, the first portable commercial sequencer of its kind in 2014 [1, 2]. *MinION*, being a small handheld device, revolutionized genomics research through its superiority in portability. The capability of nanopore sequencers to produce ultra-long reads is another advantage, especially in achieving better genome assemblies [3, 4]. Nanopore sequencers provide access to real-time current signals when a DNA strand traverses through the pore. This real-time streaming capability foster an exciting opportunity called selective sequencing [5] (also known as targeted sequencing, adaptive sampling), which has many current and promising future applications [6].

ONT’s *ReadUntil* API enables real-time bi-directional communication with an ONT sequencer: to obtain the current signals of the DNA strands being sequenced and to send back signals such as accept/reject. A rejection signal reverses the voltage difference across the pore so that the DNA strand is ejected from the pore. This can be used to quickly free a pore from sequencing an unwanted DNA strand and allow a new DNA strand to be sequenced. This can also be used to sequence selected DNA strands from a pool of different DNA species.

Successful selective sequencing is in many ways beneficial given its effective flow-cell utilization and reduced cost for sequencing. Efficient and accurate real-time analysis of the current signals streamed from the nanopore sequencer is the key to successful selective sequencing. However, existing methods used in selective sequencing to accurately compare the current signal to a target reference genome are still computationally expensive. They hence would negate the true real-time sequencing capability. Selective sequencing being relatively new, potential improvements to the performance of selective sequencing in many areas are yet to be discovered. In this paper, we investigate methods to improve the accuracy and speed of selective sequencing compared to current methodologies and present a deep neural network-based method called *DeepSelectNet*.

### Related Works

The very first attempt to realize the concept of selective sequencing used the Dynamic Time Warping (DTW) algorithm to align the raw current signal directly to a synthetic signal generated from the reference sequence [5]. However, due to the high computational complexity of DTW, this method could not scale beyond references that are longer than a few mega bases. Then, a method called RUBRIC was introduced by Edwards et al. [7], which performed real-time basecalling using Nanonet basecaller, followed by read alignment to the conventional nucleic acid references. RUBRIC delivered significant benefits in speed, scalability, and flexibility over the DTW [5] approach. Similar to RUBRIC, Readfish [8] is another basecalling based approach for selective sequencing. Readfish [8] utilizes the GPU-accelerated Guppy basecaller from ONT and Minimap2 mapper [9]. Readfish [8] is flexible and scales well to larger genomes when a powerful GPU is available.

UNCALLED [10] is a recent method that revitalises the raw signal alignment in the signal space for selective sequencing. UNCALLED probabilistically derive k-mers that can be presented in a signal and performs a comparison with a reference encoded using a FM-index [11]. Raw signals are first converted to events and the probability of each event is matched with a possible k-mer in a probabilistic k-mer model provided by ONT. UNCALLED [10] scaled much better than the DTW-based method [5], however, does not support large (>1 Gbase) or highly repetitive references.

In recent years, researchers increasingly turned their direction to deep learning approaches to improve basecalling accuracy. As the first such, a recurrent neural network approach was introduced by DeepNano [12] to predict the k-mers from raw signal segments. Then, ONT’s official basecallers followed a similar approach. According to Wick et al. [13], modern basecalling tools were able to achieve a significant improvement in basecalling with the introduction of deep learning approaches (from ~85% to 90%).

Recently, deep learning-based methods for selective sequencing started to emerge. SquiggleNet [14] is the first deep learning-based tool to classify reads based on electrical signals to perform selective sequencing. SquiggleNet adopts the convolution architecture from ResNet [15], modified to perform convolution in one dimension. It claims 90% accuracy in classifying human DNA over bacterial DNA. Also, considering the computational resources, SquiggleNet outperforms alignment-based approaches with much lesser memory consumption. However, when we benchmarked SquiggleNet across several viral and bacterial datasets, we observed that on some dataset combinations, SquiggleNet performed worse than base alignment-based methods. For instance, on datasets comprised of Corona Virus & Zymo metagenome and Corona Virus & Yeast, SquiggleNet classification accuracy could not go beyond 79%, where alignment-based approaches reached 90% (discussed under Results).

Another deep learning-based approach for selective sequencing is discussed in Danilevsky et al. [16]. The mentioned work applies deep learning models to classify mitochondrial DNA against human genomic DNA, with no reliance on labelled data to classify between the two DNA groups. However, the applicability of the mentioned method for complex inter-species classification is yet to be explored. The most recent work named *baseLess* is also a deep learning approach that relies on features of salient k-mers rather than reads as a whole, explicitly designed to work with MinION [17]. The authors have reported a performance similar to SquiggleNet, which does not surpass the performance of Guppy+Minimap2.

### Our Contribution

The existing techniques for selective sequencing have room to improve effectiveness and efficiency. We introduce DeepSelectNet, a far superior deep learning-based method, which utilizes an improved deep neural network. Also, improved techniques of raw data preprocessing enable better feature extraction with limited training data. DeepSelectNet was able to achieve over 90%(on average ~95%) classification accuracy for five datasets that we tested.

## Results

DeepSelectNet is a deep neural network based method capable of classifying species DNA directly using nanopore current signals with superior classification accuracy. DeepSelectNet is built on a convolutional architecture based on ResNet’s residual blocks [15]. Similar to SquiggleNet [14], DeepSelectNet also utilizes one-dimensional convolutional layers to perform 1D convolution over nanopore current signals in the time domain. However, DeepSelectNet relies on neural net regularization to minimise model complexity thereby reducing the overfitting of data (see Methods and Materials).

### Experiments details

Four publicly available datasets sequenced on ONT MinION/GridION, namely, SARS-CoV-2 virus (*Cov*), Zymo bacterial mixture (*Zymo*), Saccharomyces cerevisiae (*Yeast*), and Chlamydomonas algae (*Chlamy*) were used for evaluation. We prepared five combinations out of these datasets to measure the performance of DeepSelectNet: *Cov* &*Zymo*, *Cov* &*Yeast*, *Cov* &*Chlamy*, *Zymo*&*Chlamy*, and *Yeast* &*Chlamy* (detailed in Methods and Materials). For each combination, training of DeepSelectNet was done using 20000 reads from each species (thus a balanced dataset). Before training, reads were preprocessed by trimming each read’s first 1500 signal samples to eliminate the adaptor sequence (also, stall and barcodes if present; Supplementary Figure S1). This 1500 value that corresponds to ~160 bases is an overestimation for certain reads, however, it is a safe approach to eliminate unwanted portions from most of the reads [14]. Out of the remaining signal, 3000 signal samples were normalized and taken for the training. Therefore, a read to qualify for the training should have at least 4500 signal samples. The labels of the reads – from which species a given signal segment is from – were fed to the training algorithm.

### Accuracy Comparison of DeepSelectNet against existing methods

DeepSelectNet outperformed SquiggleNet in terms of test accuracy (defined in Methods and Materials) for all the dataset combinations (Fig. 1, Supplementary Table S1). The best accuracy for DeepSelectNet was observed for *Cov* &*Chlamy* (98.65%), while the best improvement compared to SquiggleNet was observed for *Cov* &*Zymo* (accuracy improvement by ~12%) as depicted in Fig. 1. For all dataset combinations, DeepSelectNet achieved a test accuracy above 90%.

**Figure 1.**
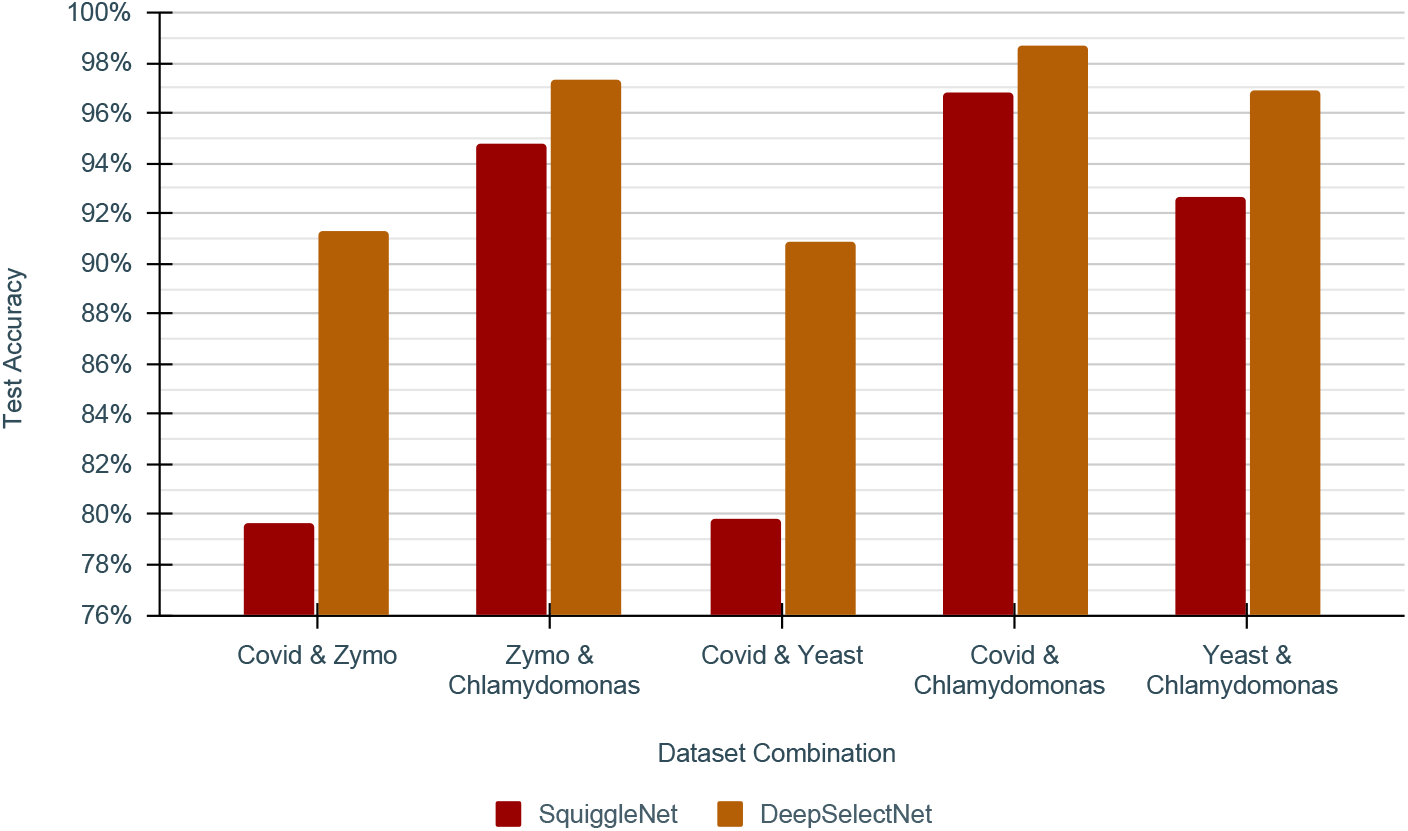
Test Accuracy comparison of SquiggleNet vs DeepSelectNet across five dataset combinations

In addition to the comparison with SquiggleNet, DeepSelectNet’s performance was then compared against: Guppy high accuracy basecalling on complete reads followed by Minimap2 [9] mapping (*baseline*); Guppy high accuracy basecalling on the first 4500 signal samples of each read^[1]^ followed by Minimap2 mapping (*Guppy_hac+Minimap2*); and, Guppy fast basecalling on the first 4500 signal samples of each read followed by Minimap2 mapping (*Guppy_fast+Minimap2*). Accuracy results are shown in Fig. 2. The *baseline* (Fig. 2) that uses the complete read (impractical for selective sequencing) was to get an estimate of the upper margin accuracy a certain computational method could potentially achieve. *Guppy_hac+Minimap2* mirrors a real-time scenario as in DeepSelectNet’s intended use, where the whole read is not available for classification. *Guppy_fast+Minimap2* follows the approach used in Readfish [8]. While we expected the accuracy of DeepSelectNet to be lower than the baseline, interestingly, DeepSelectNet outperformed the baseline method in three (*Zymo*&*Chlamy*, *Cov* &*Chlamy* and *Yeast* &*Chlamy*) out of five dataset combinations (Fig. 2, Supplementary Table S2). Compared to *Guppy_hac+Minimap2* that considers the realtime scenario (unlike the baseline method that uses complete reads), DeepSelectNet was performing better in four out of five dataset combinations (except *Cov* &*Zymo*; Fig. 2). *Guppy_fast+Minimap2* accuracy was somewhat in close proximity to *Guppy_hac+Minimap2*, yet lower than DeepSelectNet in all dataset combinations. Also, it was observed that *Guppy_fast+Minimap2* has the worst performance in three out of five dataset combinations. As the predecessor of DeepSelectNet, the SquiggleNet performed as the second best method in three out of five dataset combinations, when considering real-time scenarios.

**Figure 2.**
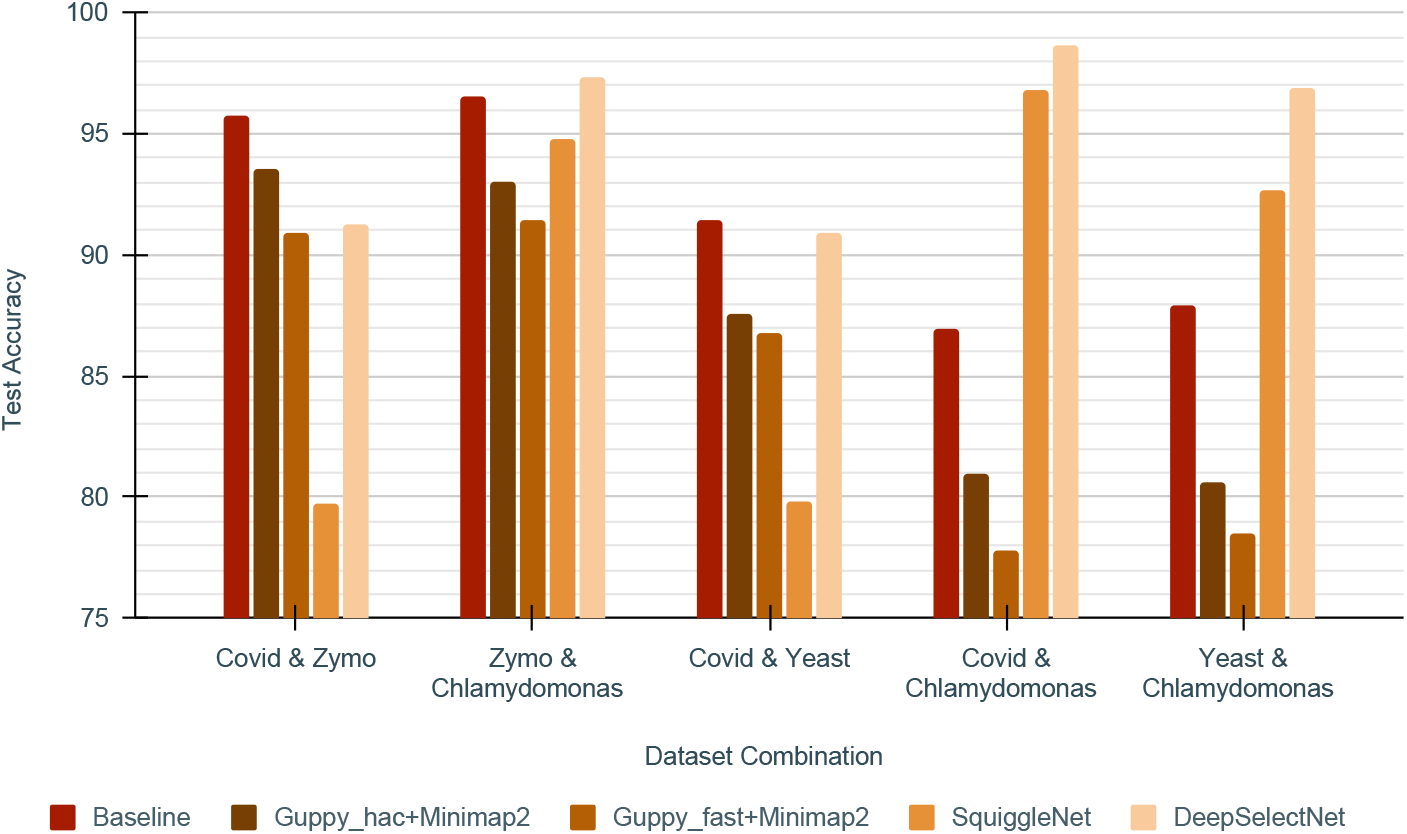
Test Accuracy comparison of DeepSelectNet against existing methods across five dataset combinations

### Runtime performance comparison

In addition to accuracy benchmarking, we also benchmarked the execution time for inference (see Fig. 3 where the per-read execution time is plotted, Supplementary Table S3). A setup similar to that discussed earlier (previous subsection) was used for these experiments. Among the methods experimented with, *Guppy_fast+Minimap2* had the best execution time throughout all datasets.

**Figure 3.**
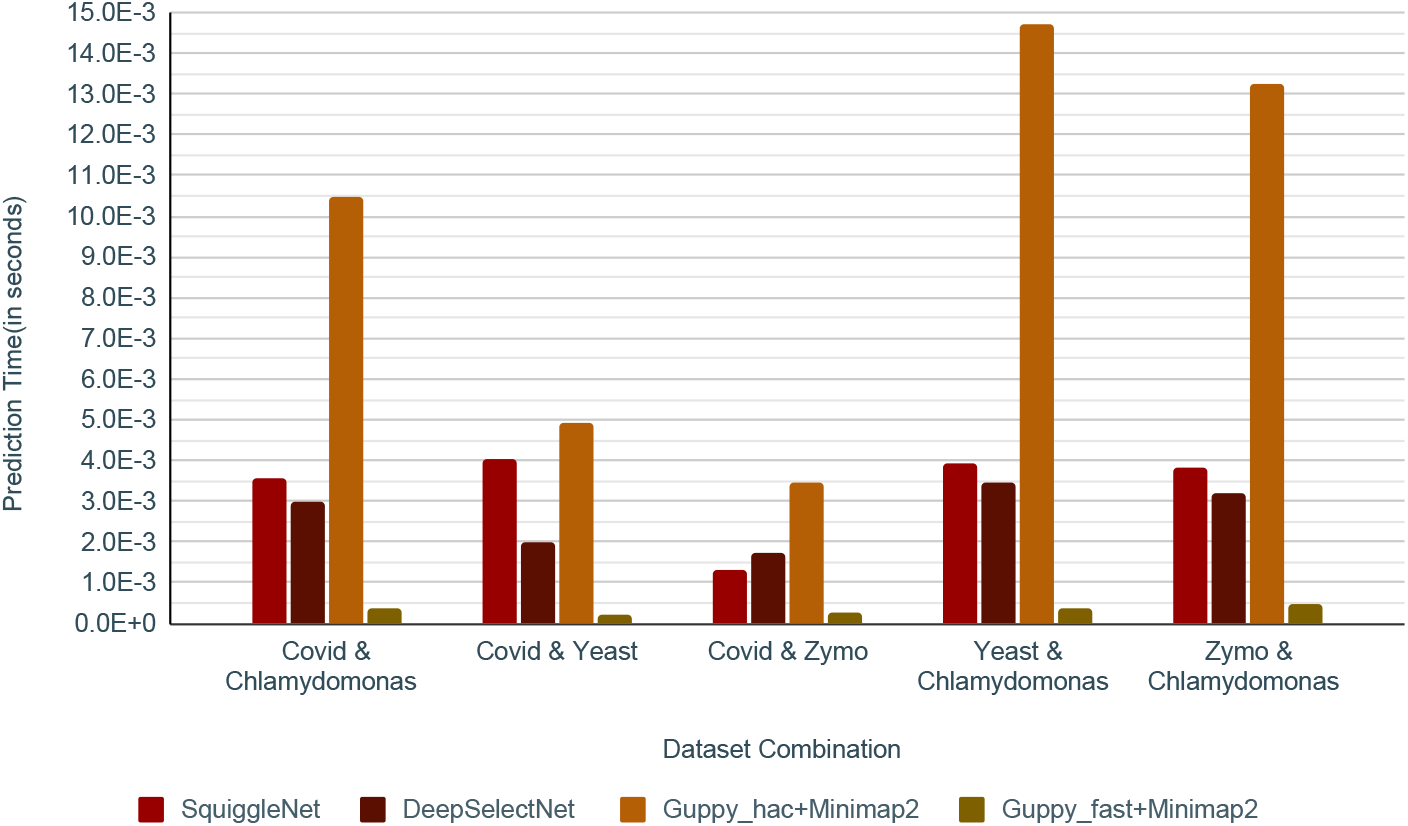
Inference runtime comparison of DeepSelectNet against other methods across five dataset combinations

*Guppy_hac+Minimap2* was the slowest (Fig. 3). This drastic difference can be accounted for by the guppy configuration of the model being used in the two methods. Compared to SquiggleNet, DeepSelectNet has the best runtime performance in four out of five datasets, except *Cov* &*Zymo* where SquiggleNet was faster compared to DeepSelectNet. Even though DeepSelectNet is not as fast as *Guppy_fast+Minimap2*, DeepSelectNet’s superior accuracy must be noted. Also, note that Guppy which is the ONT’s production basecaller written in C/C++ is likely to be well-optimised for execution performance, unlike DeepSelectNet which was a prototype developed using Python. While it is not in the scope of this work to optimise DeepSelectNet for execution performance, it is logical to believe that reimplementing using C/C++ and performing optimisations could lead to improved execution time.

### Precision, recall, and F1 scores of DeepSelectNet

Similar to Accuracy, DeepSelectNet shows scores >89% on other performance metrics: precision, recall, and F1 as depicted in Fig. 4 (Supplementary Table S4). Given all the datasets used for training and testing are balanced as explained earlier, being able to observe both high precision and recall demonstrates that the classification is not biased^[2]^ towards a certain species in a dataset combination while also having a low variance^[3]^ of predictions.

**Figure 4.**
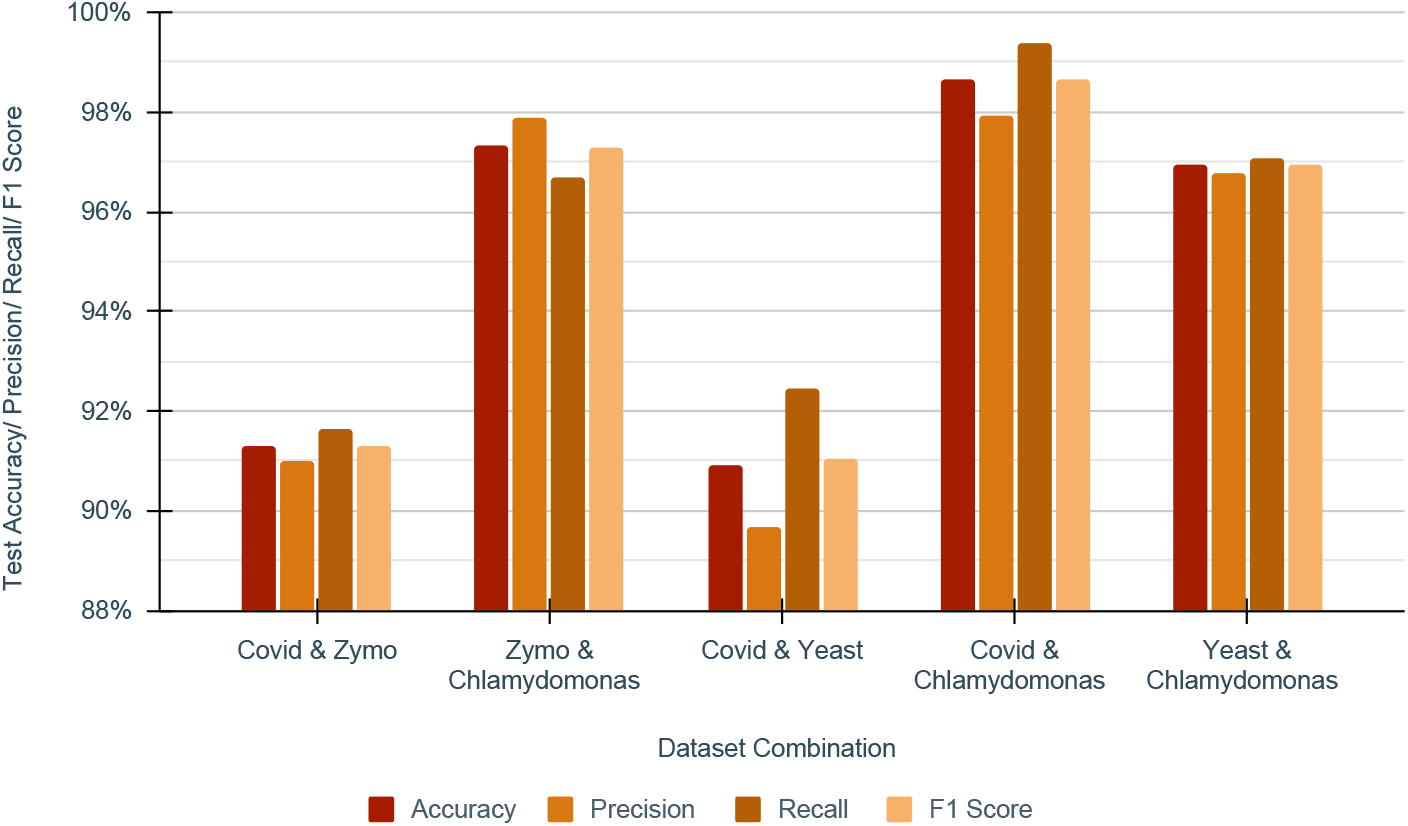
DeepSelectNet’s performance metrics across five dataset combinations

## Discussion

DeepSelectNet shows promising improvements in accuracy compared to existing deep learning-based and alignment-based approaches for selective sequencing. DeepSelectNet is a result of observing that the existing deep learning-based methods, such as SquiggleNet, had room to improve further. The model’s ability to classify at higher accuracy, precision and recall demonstrates that the classification is unbiased to a particular species.

### Impact of number of reads and segment size on accuracy

We measured the model accuracy while increasing the number of reads used for training. We observed that the number of reads used for training improves the classification accuracy (Fig. 5). It is likely that the increase in accuracy is due to the increased genome coverage when the number of reads is increased (coverage is defined in Methods and Materials). When the coverage of the genome by a dataset is high, the model will have more features to learn about the species (and also the effect due to noise is diminished) and classify them better against the other species.

**Figure 5.**
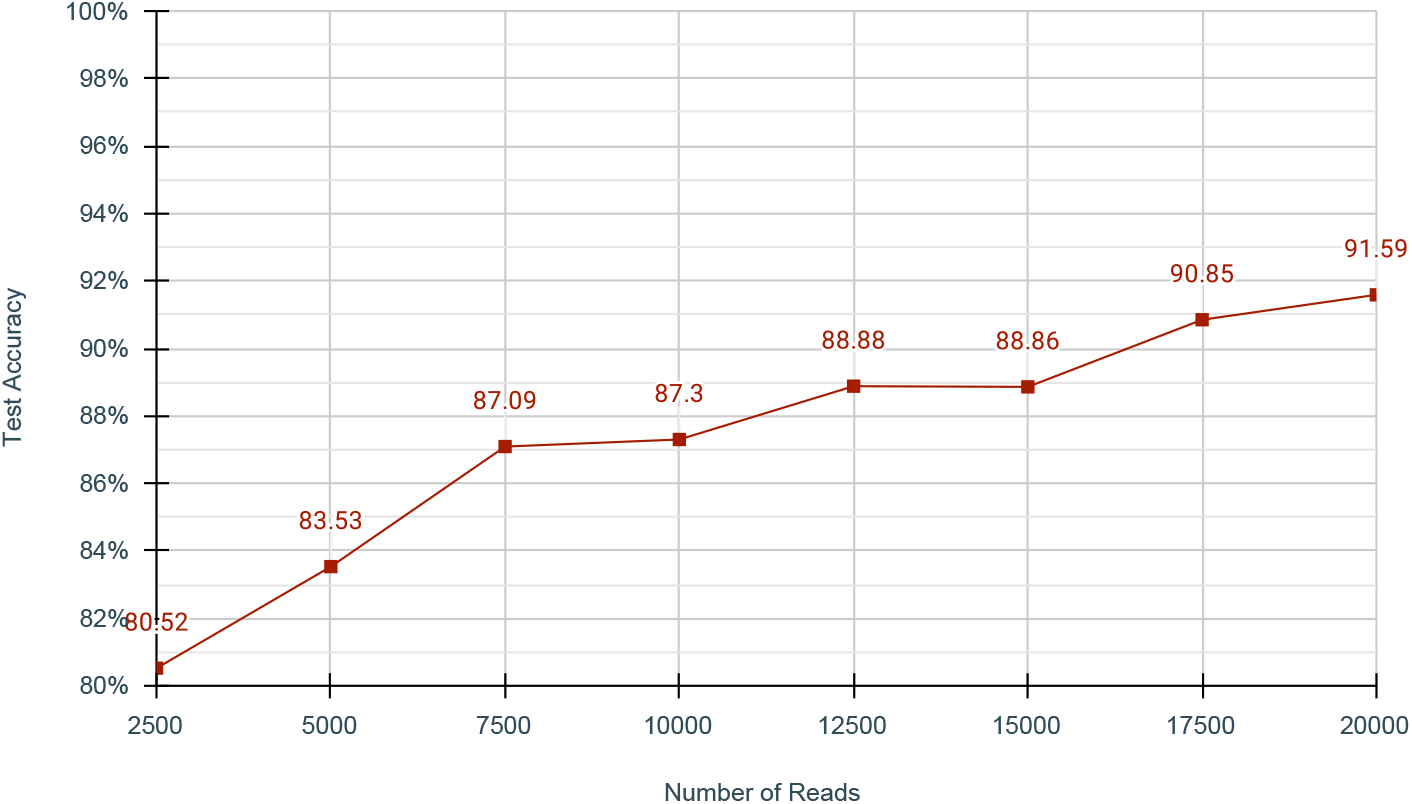
Accuracy of DeepSelectNet against varying number of reads for Cov&Zymo dataset. The segment size (excluding the 1500 samples trimmed at the beginning) was kept constant at 3000 samples. Refer Supplementary Table 14 for details Genome Coverage in each case.

We also measured the model accuracy against different signal segment lengths (no. of signal samples in each read) and noticed that the accuracy improves with longer segments (Fig. 6). However, the improvement becomes less significant after ~4000 signal samples (Fig. 6). We selected a segment size of 3000 signal samples for our experiments above (~330 bases), considering the time to train, real-time inference requirement and fair comparison with SquiggleNet [14] (SquiggleNet also used 3000 as the segment length). A longer signal segment means having a larger k-mer size that allows distinguishing species better, thereby the model may classify better with longer segments. Having a longer segment increases the genome coverage, which also may improve the model performance as discussed before.

**Figure 6.**
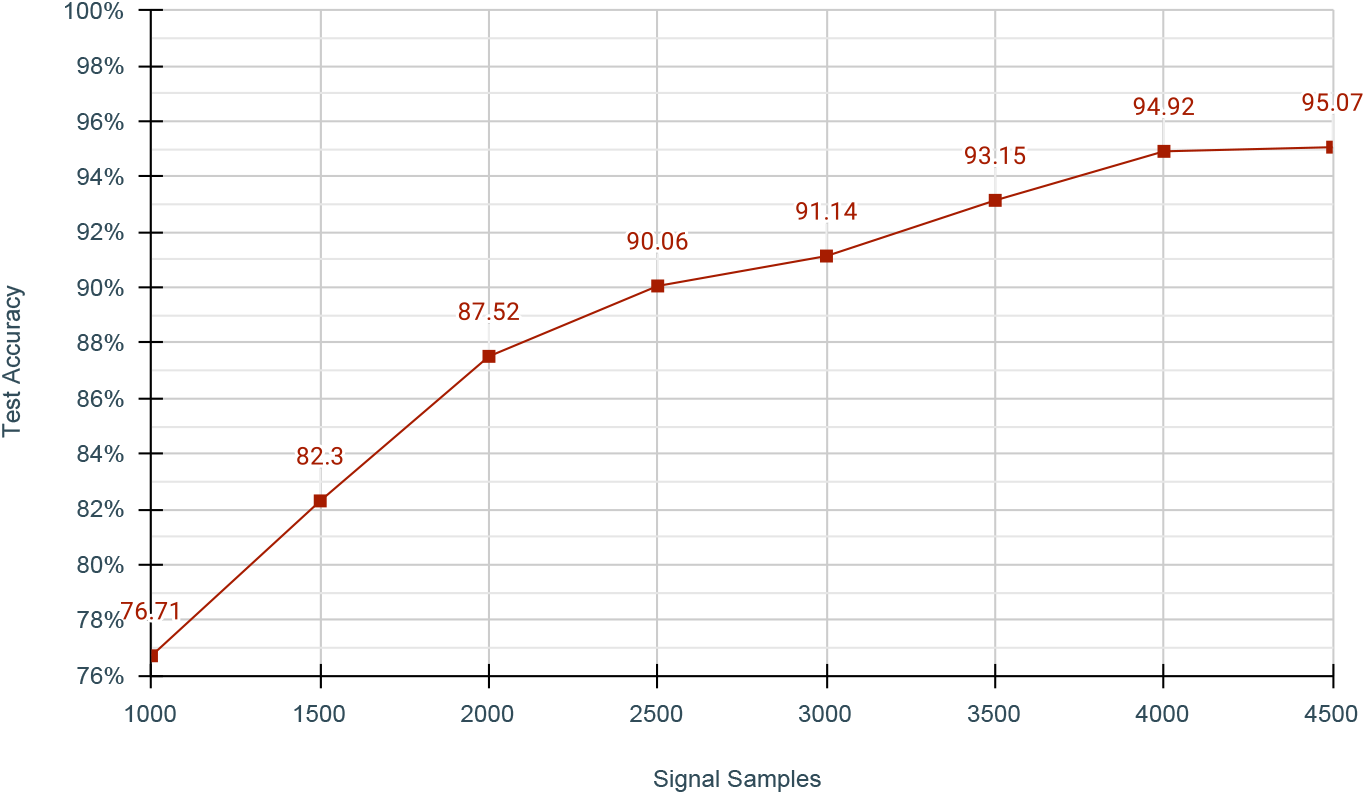
Accuracy of DeepSelectNet agianst the signal segment length for Cov&Zymo dataset. The number of reads used for training was kept constant at 20,000. Note that the segment length excludes the first 1500 signal samples that are trimmed. Refer Supplementary Table 13 for details Genome Coverage in each case

### DeepSelectNet with human datasets

Inter-species experiments we performed with *Human*&*Zymo* and *Human*&*Covid*, produced 92.26% and 96.27% test accuracies, respectively (Supplementary Table S9). Human data were from the publicly available NA12878 reference Genome [18]. For these experiments, 20,000 reads from each species were used (relates to 0.008×, 400×, and 0.4× genome coverage for human, covid and zymo respectively, Methods and Materials, Supplementary Table S12). Interestingly, even without seeing the complete genome (coverage <0.008× for human), the model could classify accurately. This could be because, even though the coverage is less, still the features are sufficiently different (an analogy would be k-mers distribution between humans and bacteria/viruses being different). However, it also raised a concern if the model is picking some other difference amongst the datasets than the species (for instance, differences in library preparation method and/or minor differences in the flowcells in the publicly available datasets sequenced by different labs at different times). To eliminate this concern, we simulated synthetic nanopore raw signal data for the two species Yeast and Chlamydomonas using *Squigulator* [19] that relies on a pore-model indicating the expected current level for each k-mer. When this simulated data was used for training and testing, DeepSelectNet could achieve a test accuracy of 97.10% (Supplementary Table S11). Therefore, this experiment demonstrates that DeepSelectNet actually classifies based on features of species, rather than being misled by any other non-species related difference.

Though the DeepSelectNet performed great for inter-species classification as explained throughout the paper (accuracy ~95% on average), its performance was not as great for intra-species classification (that is, classifying reads based on a genomic region of the same species). For example, when chromosome 21 and chromosome 22 from the human genome (publicly available NA12878 data available under [20]) were used to train DeepSelectNet (chr21 as the positive dataset and chr22 as the negative dataset), the training accuracy was below 60% in all cross folds. Increasing the number of reads for each chromosome from 20,000 to 100,000 (signal segment length of 3000 samples), improved the training accuracy by a small margin up to 64%. With 35,000 reads from each chromosome at a signal segment length of 10,000 (~1100 bases), the training accuracy improved to 73%. It is possible that DeepSelectNet is reaching its performance limitations to learn fine features among different chromosomes or different regions of a species. Unfortunately, our computational infrastructure restricted us from exploring further on larger datasets than this.

## Methods and Materials

### DeepSelectNet Model Architecture

The first layer in DeepSelectNet is a one-dimensional Convolutional Neural Network (CNN) with 20 channels. The rest of the residual blocks (Similar to ResNet) are implemented additionally with dropout layers [21] to effectively regularize any overfitting effects that the layer complexity might introduce. This helps to mitigate the effect of the model picking up on statistical noise in the training data, which could result in poor performance when the model is evaluated on unseen data. During training, 10% layer outputs are randomly ignored so that the layer will be treated like a layer with a different number of nodes and edges than the layer before. Therefore, each update to a layer during training is performed differently in comparison to the updated layer. The four residual layers, which include two bottleneck blocks, are stacked together following a mean pooling layer and lastly, a fully-connected layer activated with a sigmoid function that flattens the output for classification.

### DeepSelectNet Implementation

DeepSelectNet was developed in Python programming language using the Keras framework^[4]^. Core functionalities of DeepSelectNet are distributed among three scripts:

- *preproccesor.py* that pre-processes datasets prior to training (read filtering, signal trimming, signal segmentation, signal normalization and splitting to batches).
- *trainer.py* that trains the pre-processed data (change hyperparameters of the model including classifier network, loss function, train to validation split ratio, number of cross folds, epochs, data batch size, etc.).
- *inference.py* that inferences test data using a trained model (data preprocessing similar to *preproccessor.py* is done in this script before the inference).

### Data Pre-processing

Four publicly available datasets sequenced on ONT MinION/GridION were used for the experiments (Table 1). These datasets contained raw signal data in single-FAST5 format (one file per each read), which we converted to SLOW5 format using slow5tools [20] to enable convenient and efficient file manipulation. Then, 40,000 reads containing at least 4500 signal samples were extracted from each dataset. Using these extracted reads, five dataset combinations were created, namely *Cov* &*Zymo*, *Cov* &*Chlamy*, *Cov* &*Yeast*, *Yeast* &*Chlamy* and *Zymo*&*Chlamy*. Each dataset combination was partitioned equally for training and testing. For instance, the *Cov* &*Yeast* dataset combination which contains 40,000 from *Cov* and 40,000 from *Yeast* is partitioned such that 20,000 *Cov* reads and 20,000 *Yeast* reads are for training, and the rest for testing. Thus, the created datasets were balanced datasets where a species’ contribution to each dataset combination was equal. Note that this count of 20000 reads was a value that we empirically determined to adequately provide genome-wide coverage for these dataset combinations (See the Discussion for more information).

**Table 1.** Datasets and their sources

| Species | Dataset | Source |
| --- | --- | --- |
| SARS-CoV-2 | Cov | https://community.artic.network/t/links-to-raw-fast5-fastq-data-for-artic-protocol/17 |
| Zymo Metagenome | Zymo | https://github.com/LomanLab/mockcommunity |
| Chlamydomonas | Chlamy | https://sra-download.ncbi.nlm.nih.gov/traces/era20/ERZ/003237/ERR3237140/Chlamydomonas_0.tar.gz |
| Saccharomyces cerevisiae | Yeast | https://www.ncbi.nlm.nih.gov/bioproject/PRJNA510813 |

Each read from the dataset is first trimmed by removing the initial 1500 signal samples. As discussed earlier, trimming is essential to get rid of the read adaptor (and barcode if exists, Supplementary Figure S1). Then, the raw signal samples (16-bit integers) are converted to pico-amperes using the equation: 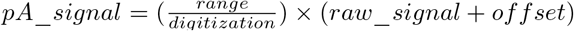. Next, four random segments (where each segment is 3000 signal samples) are obtained from this converted pico-ampere signal^[5]^. Taking multiple signal segments from a single long read provides more feature space than taking a single segment and directly affects on model performance. This technique helped DeepSelectNet to increase its accuracy to a great extent (Supplementary Table S10). Value four as the number of segments was derived empirically, considering the model accuracy, the average length of the reads and the time to train.

Each such segment is then normalized with modified version of z-score that uses median absolute deviation (MAD): 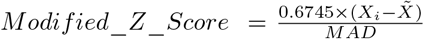; where 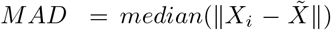 and 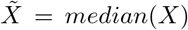. The median absolute deviation used in the modified z-score is tolerant to extreme outliers (Supplementary Figure S2) present in the data. The threshold used for MAD in data preprocessing had an effect on model training (Fig. 7, Supplementary Table S7). The MAD threshold affects the number of outliers being filtered, producing a differently performing model. Therefore, the better the outliers are filtered, the better the model trains. The best MAD threshold could depend on many factors such as the species, sequencing device and sample purity and thus we determined this threshold empirically (Supplementary Figure S3). For this, We conducted experiments with MAD = 3, 5, and 10 as the thresholds and picked the value that produced the best accuracy for a given dataset combination (Fig. 7).

**Figure 7.**
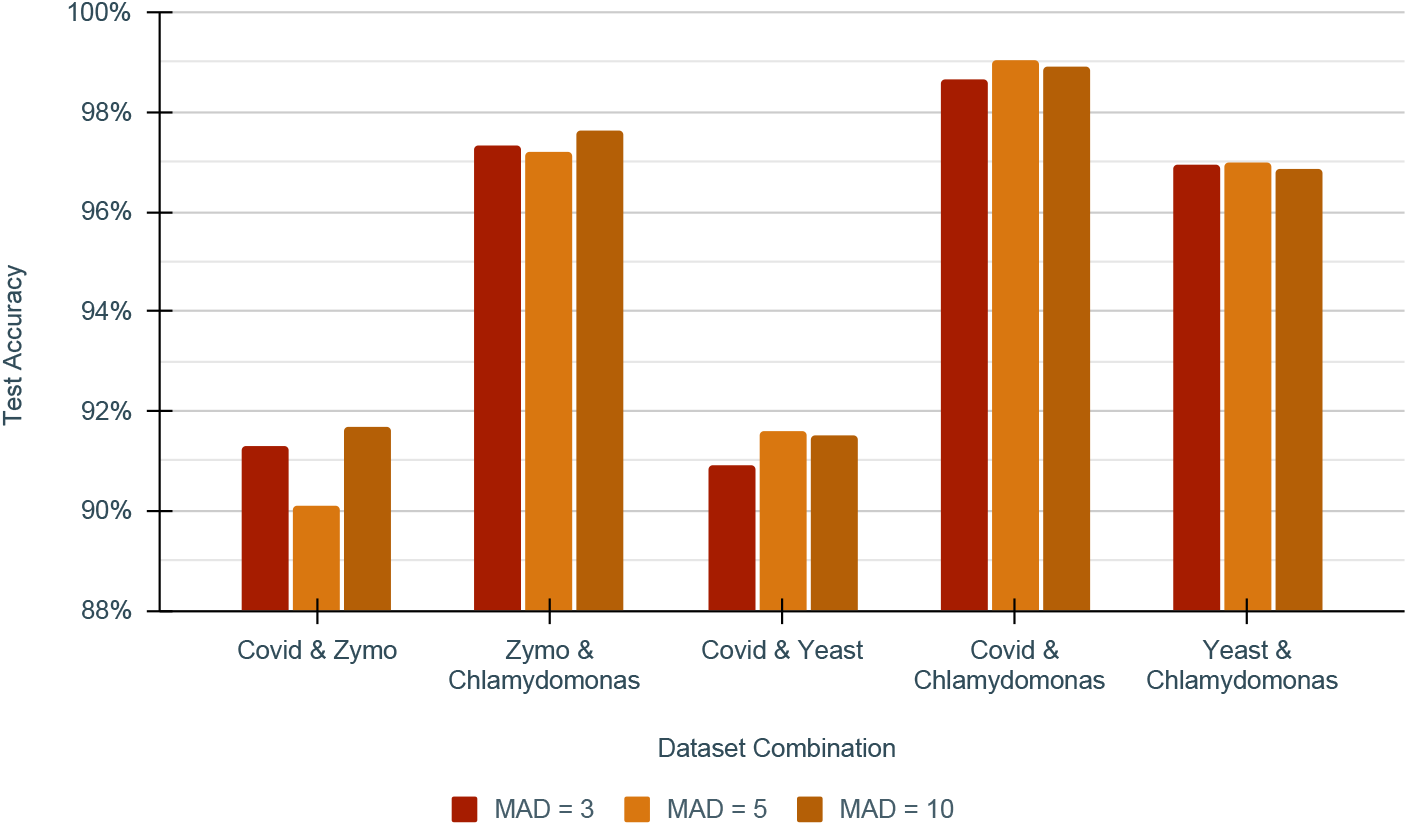
Impact of Median Absolute Deviations(MAD) on DeepSelectNet’s Test Accuracy

### Training

In a given dataset combination, the training data partition (explained above) was again partitioned in 7:3 ratio, where 70% of training data was used for training and the rest for validation. As an example, *Cov* &*Yeast* dataset combination containing 20,000 training reads from *Cov* and 20,000 training reads from *Yeast* is partitioned such that 14,000 *Cov* reads and 14,000 *Yeast* reads are for training, and the rest for validation.

Additionally, cross-fold validation was used to generalise training to ensure a minimally biased model. The training accuracy of trained models in each of the 5 folds lies in close proximity (Fig. 8, Supplementary Table S8), demonstrating that training has generalised.

**Figure 8.**
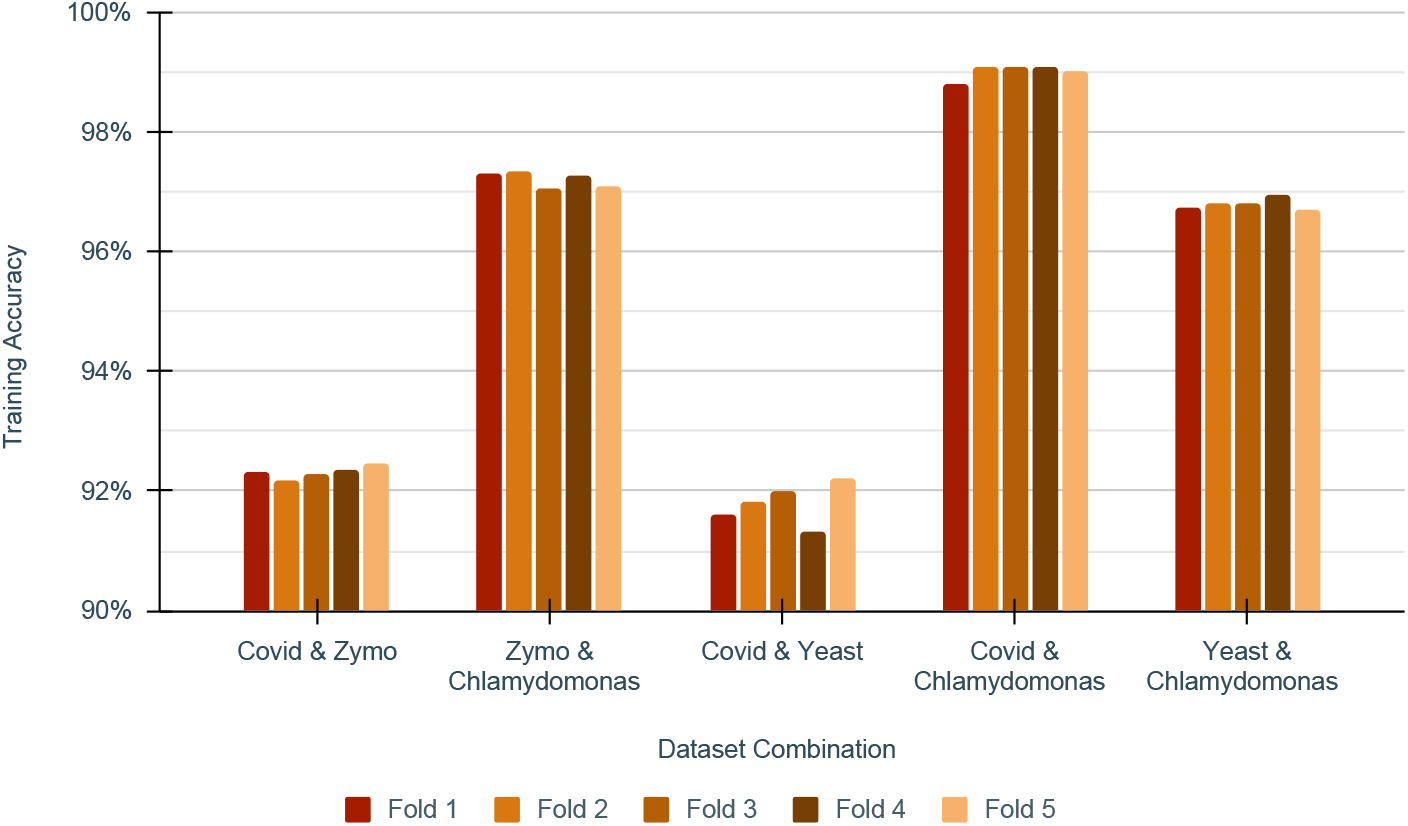
Training Accuracy of DeepSelectNet’s across five cross folds

### Evaluation

The best performing model of the five training cross folds was selected for the evaluation. All evaluations were done with previously unseen test data created in the preprocessing step. The Test data is also first preprocessed: the first 1500 signal samples in each read are trimmed off; the next 3000 signal samples are taken (note that only the first segment that represents the beginning of the read is taken for testing; we do not take 4 segments as in training); converted to pA and normalized using the modified z-score as done for training.

The inference was performed in batches of 1000 reads to ensure higher GPU occupancy and also indicates a scenario where reads are being sequenced in parallel on a nanopore sequencer with 1000 channels. The model training and the accuracy-related benchmarks were conducted on a system powered by an Intel Xeon CPU E5-1630 v3 @ 3.70GHz CPU, 32 GB of RAM, 2TB of HDD storage, and an Nvidia Quadro K40 GPU. As this was a shared workstation unsuitable for reliable timing measurements, runtime benchmarks were performed on another dedicated workstation powered by an Intel Xeon Silver 4114 CPU @ 2.20GHz CPU, 384 GB RAM, 6TB NVME SSD storage and a Tesla V100-16G GPU.

### Modifications to SquiggleNet for comparison against DeepSelectNet

We modified SquiggleNet to accept SLOW5 file format [20] as an input (SquiggleNet only accepted single-FAST5 files and manipulation of those files is inefficient and inconvenient). This modification does not affect the model classification accuracy as the underlying raw signal data is identical despite the file format used. The modified source code of SquiggleNet is available in the link provided under Availability of data and materials.

### Baseline, Guppy_hac+Minimap2 and Guppy_fast+Minimap2

The *baseline* approach uses complete reads whereas *Guppy_hac+Minimap2* and *Guppy_fast+Minimap2* uses only the first 4500 signal samples in a read (see Results). Guppy version 6.1.3 and Minimap2 version 2.20 were used for the experiments. For *baseline* and *Guppy_hac+Minimap2*, the high-accuracy basecalling mode *dna_r9.4.1_450bps_hac* was used for Guppy and Minimap2 was executed with base alignment enables (with -c option). For *Guppy_fast+Minimap2*, fast base-calling mode was used with Guppy and Minimap2 was executed without base alignment to mirror the approach used in Readfish [8].

### Genome Coverage computation

Genome Coverage denotes the ratio between the number of bases appearing in the given dataset to the actual number of bases in its reference genome. The formula we used to compute the genome coverage seen by the model during training for a given dataset is given below.

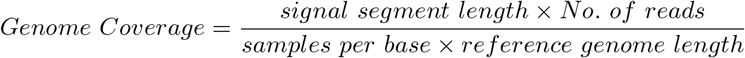

The signal segment length is the number of samples in each signal segment used for training (excluding the samples trimmed at the beginning of the read). Samples per base refer to the number of signal samples that correspond to a single base, which is determined by the translocation speed of the DNA strand through the pore and the sampling rate of data acquisition. For DNA on R9.4.1, 450 bases/s at 4000 Hz gives ~8-10 Samples per base.

### Accuracy Metrics

Accuracy metrics and related terms used for evaluating the classification performance of the models in this work are defined below.

An outcome where the model predicts a read as positive when the read is actually from the targeted (positive) dataset/species is considered a *True Positive (TP)*. An outcome where the model predicts a read as negative when the read is actually from the non-targeted (negative) dataset/species is considered a *True Negative (TN)*. An outcome where the model predicts a read as positive when the read is actually from the non-targeted (negative) dataset/species is considered a *False Positive (FP)*. An outcome where the model predicts read as negative when the read is actually from the targeted (positive) dataset/species is considered a *False Negative (FN)*. Accuracy that denotes the overall accuracy of the model with correct predictions over total predictions is computed as 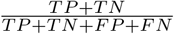; precision is computed as 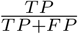; Recall is computed as 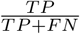; and F1 score 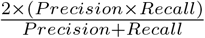. However, in experiments from baseline method, Guppy_hac+Minimap2 and Guppy_fast+Minimap2, the unmapped reads accounted neither as false positives nor false negatives. Therefore in such cases, Precision, Recall and F1 Score are approximations with available prediction statistics.

## Conclusion

Nanopore selective sequencing is an emerging area of genome sequencing that is gaining popularity due to its efficiency and low cost. There have been several attempts to realize selective sequencing with better accuracy and speed. In this paper, we presented DeepSelectNet, a deep learning-based approach for species classification using selective sequencing that outperforms existing methods. Amongst five inter-species datasets evaluated, DeepSelectNet achieved >90% accuracy (on average ~95%) for all datasets. DeepSelectNet was bench-marked against existing methods (both deep-learning and non-deep-learning methods) and produced exceedingly top performances across the vast majority of the datasets. In some cases, DeepSelectNet on partial reads (4500 signal samples at the beginning of each read) outperformed the alignment-based approach (high accuracy basecalling + alignment) done on complete reads. DeepSelectNet also performed well in classifying human DNA from bacteria/viral data (>90% accuracy). However, DeepSelectNet could only achieve <70% for intra-species classification such as classifying based on different regions of the same species. Thus, the architecture of DeepSelectNet is more suitable for inter-species classification. In terms of execution time, DeepSelectNet outperformed the existing deep-learning-based method. However, DeepSelectNet being a Python-based prototype was several times slower than the C/C++-based Guppy (fast basecalling) + Minimap2 combination. Re-implementation of DeepSelectNet in C/C++ followed by performance tuning would potentially achieve comparable runtime performance.

## Availability of data and materials

The source code for DeepSelectNet is available on Github at https://github.com/AnjanaSenanayake/DeepSelectNet. The source code for modified SquiggleNet is available on Github at https://github.com/AnjanaSenanayake/SquiggleNet/tree/slow5-support. All instructions for using the tool can be found in Supplementary Note 1.

Datasets used are publicly available datasets and their links can be found in Table 1. Additionally, a curated dataset used for DeepSelectNet experiments can be found at https://doi.org/10.5281/zenodo.7111366

## Footnotes

[1] translates to ~300 bases when the adaptor and barcode are removed

[2] The difference between the average prediction of the model and the actual target - high bias means under-fitting

[3] The variability between the actual target and predicted target - high variance means over-fitting.

[4] Keras is a high-level API built up on Tensorflow to develop deep neural networks

[5] there could be overlaps when randomly taking these segments; if the remaining signal length is only 3000 samples, all the four segments will be identical

