## Supplementary Information for "DeepSelectNet: Deep Neural Network Based Selective Sequencing for Oxford Nanopore Sequencing"

#### Table S1. Test accuracy comparison of SquiggleNet vs DeepSelectNet across five dataset combinations

| Dataset | Squigglenet | DeepSelecNet |
|---|---|---|
| **Covid & Zymo** | 79.69% | 91.28% |
| **Zymo & Chlamydomonas** | 94.81% | 97.31% |
| **Covid & Yeast** | 79.83% | 90.9% |
| **Covid & Chlamydomonas** | 96.82% | 98.65% |
| **Yeast & Chlamydomonas** | 92.69% | 96.93% |

#### Table S2. Test accuracy comparison of DeepSelectNet against existing methods across five dataset combinations

| Datasets | Baseline | Guppy_hac+ Minimap2 | SquiggleNet | DeepSelectNet | Guppy_fast+ Minimap2 |
|---|---|---|---|---|---|
| **Covid & Zymo** | 95.76% | 93.53% | 79.69% | 91.28% | 90.95% |
| **Zymo & Chlamydomonas** | 96.58% | 93.03% | 94.81% | 97.31% | 91.46% |
| **Covid & Yeast** | 91.4% | 87.53% | 79.83% | 90.9% | 86.79% |
| **Covid & Chlamydomonas** | 86.94% | 80.96% | 96.82% | 98.65% | 77.83% |
| **Yeast & Chlamydomonas** | 87.94% | 80.62% | 92.69% | 96.93% | 78.49% |

#### Table S3. Prediction runtime comparison of DeepSelectNet against other methods across five dataset combinations

| Dataset | SquiggleNet | DeepSelectNet | Guppy_hac+ Minimap2 | Guppy_fast+Minimap2 |
|---|---|---|---|---|
| **Covid & Chlamydomonas** | 0.00355 ms | 0.003 ms | 0.0105 ms | 0.000375 ms |
| **Covid & Yeast** | 0.004025 ms | 0.001975 ms | 0.004925 ms | 0.0002 ms |
| **Covid & Zymo** | 0.001325 ms | 0.00175 ms | 0.00345 ms | 0.000275 ms |
| **Yeast & Chlamydomonas** | 0.00395 ms | 0.00345 ms | 0.014725 ms | 0.000375 ms |
| **Zymo & Chlamydomonas** | 0.00385 ms | 0.003175 ms | 0.013225 ms | 0.00045 ms |

**Table S4. DeepSelectNet's overall performance across five dataset combinations**

| Dataset | Accuracy | Precision | Recall | F1 Score |
|---|---|---|---|---|
| **Covid & Zymo** | 91.28% | 91% | 91.62% | 91.31% |
| **Zymo & Chlamydomonas** | 97.31% | 97.9% | 96.69% | 97.29% |
| **Covid & Yeast** | 90.9% | 89.67% | 92.45% | 91.04% |
| **Covid & Chlamydomonas** | 98.65% | 97.93% | 99.39% | 98.65% |
| **Yeast & Chlamydomonas** | 96.93% | 96.78% | 97.08% | 96.93% |

**Table S5. DeepSelectNet's performance across different read lengths in the Cov&Zymo dataset**

| Read Length | Accuracy |
|---|---|
| 1000 | 76.71% |
| 1500 | 82.3% |
| 2000 | 87.52% |
| 2500 | 90.06% |
| 3000 | 91.14% |
| 3500 | 93.15% |
| 4000 | 94.92% |
| 4500 | 95.07% |

**Table S6. DeepSelectNet's performance across different number of read samples in Cov&Zymo dataset**

| Number of Reads | Accuracy |
|---|---|
| 2500 | 80.52% |
| 5000 | 83.53% |
| 7500 | 87.09% |
| 10000 | 87.3% |
| 12500 | 88.88% |
| 15000 | 88.86% |
| 17500 | 90.85% |
| 20000 | 91.59% |

**Table S7. Impact of Median Absolute Deviations(MAD) on DeepSelectNet's accuracy**

| Dataset | MAD = 3 | MAD = 5 | MAD = 10 |
|---|---|---|---|
| **Covid & Zymo** | 91.28% | 90.09% | 91.68% |
| **Zymo & Chlamydomonas** | 97.31% | 97.22% | 97.62% |
| **Covid & Yeast** | 90.9% | 91.58% | 91.5% |
| **Covid & Chlamydomonas** | 98.65% | 99.06% | 98.92% |
| **Yeast & Chlamydomonas** | 96.93% | 96.97% | 96.87% |

**Table S8. Training Accuracy of DeepSelectNet's across five cross folds**

| Dataset | Fold 1 | Fold 2 | Fold 3 | Fold 4 | Fold 5 |
|---|---|---|---|---|---|
| **Covid & Zymo** | 92.31% | 92.19% | 92.29% | 92.34% | 92.48% |
| **Zymo & Chlamydomonas** | 97.3% | 97.35% | 97.05% | 97.26% | 97.11% |
| **Covid & Yeast** | 91.62% | 91.81% | 91.99% | 91.32% | 92.2% |
| **Covid & Chlamydomonas** | 98.82% | 99.1% | 99.11% | 99.09% | 99.01% |
| **Yeast & Chlamydomonas** | 96.73% | 96.81% | 96.8% | 96.97% | 96.7% |

**Table S9. Performance of DeepSelectNet's across dataset combinations including Human**

| Dataset | Accuracy | Precision | Recall | F1 Score |
|---|---|---|---|---|
| **Human& Covid** | 96.27% | 96.19% | 96.35% | 96.27% |
| **Human & Zymo** | 92.26% | 93.24% | 91.13% | 92.17% |
| **Human & Yeast** | 81.82% | 83.94% | 78.68% | 81.23% |
| **Human & Chlamydomonas** | 91.19% | 89.24% | 93.68% | 91.41% |

**Table S10. Accuracies of DeepSelectNet's before and after introducing segment sampling**

| | DeepSelectNet Accuracy | |
|---|---|---|
| **Dataset** | **Before** | **After** |
| **Covid & Zymo** | 84.19% | 91.28% |
| **Zymo & Chlamydomonas** | 95.54% | 97.31% |
| **Covid & Yeast** | 82.88% | 90.90% |
| **Covid & Chlamydomonas** | 97.68% | 98.65% |
| **Yeast & Chlamydomonas** | 94.48% | 96.93% |

**Table S11. DeepSelectNet performance for artificially generated signals for Yeast & Chlamydomonas**

| Species | Accuracy |
|---|---|
| Yeast | 97.39% |
| Chlamydomonas | 96.82% |
| Yeast & Chlamydomonas | 97.10% |

**Table S12. Genome Coverage of the individual species used in dataset combinations**

| Dataset | Genome Coverage |
|---|---|
| Chlamydomonas | 0.22 |
| Covid | 800 |
| Yeast | 0.67 |
| Zymo | 0.4 |

**Table S13. Genome Coverage of Covid & Zymo for different read lengths**

| | Genome Coverage | |
|---|---|---|
| Read Length | Covid | Zymo |
| 1000 | 266.6666667 | 0.1290322581 |
| 1500 | 400 | 0.1935483871 |
| 2000 | 533.3333333 | 0.2580645161 |
| 2500 | 666.6666667 | 0.3225806452 |
| 3000 | 800 | 0.3870967742 |
| 3500 | 933.3333333 | 0.4516129032 |
| 4000 | 1066.666667 | 0.5161290323 |
| 4500 | 1200 | 0.5806451613 |

**Table S14. Genome Coverage of Covid & Zymo for different number of reads**

| | Genome Coverage | |
|---|---|---|
| Number of Reads | Covid | Zymo |
| 2500 | 666.6666667 | 0.3225806452 |
| 5000 | 1333.333333 | 0.6451612903 |
| 7500 | 2000 | 0.9677419355 |
| 10000 | 2666.666667 | 1.290322581 |
| 12500 | 3333.333333 | 1.612903226 |
| 15000 | 4000 | 1.935483871 |
| 17500 | 4666.666667 | 2.258064516 |
| 20000 | 5333.333333 | 2.580645161 |

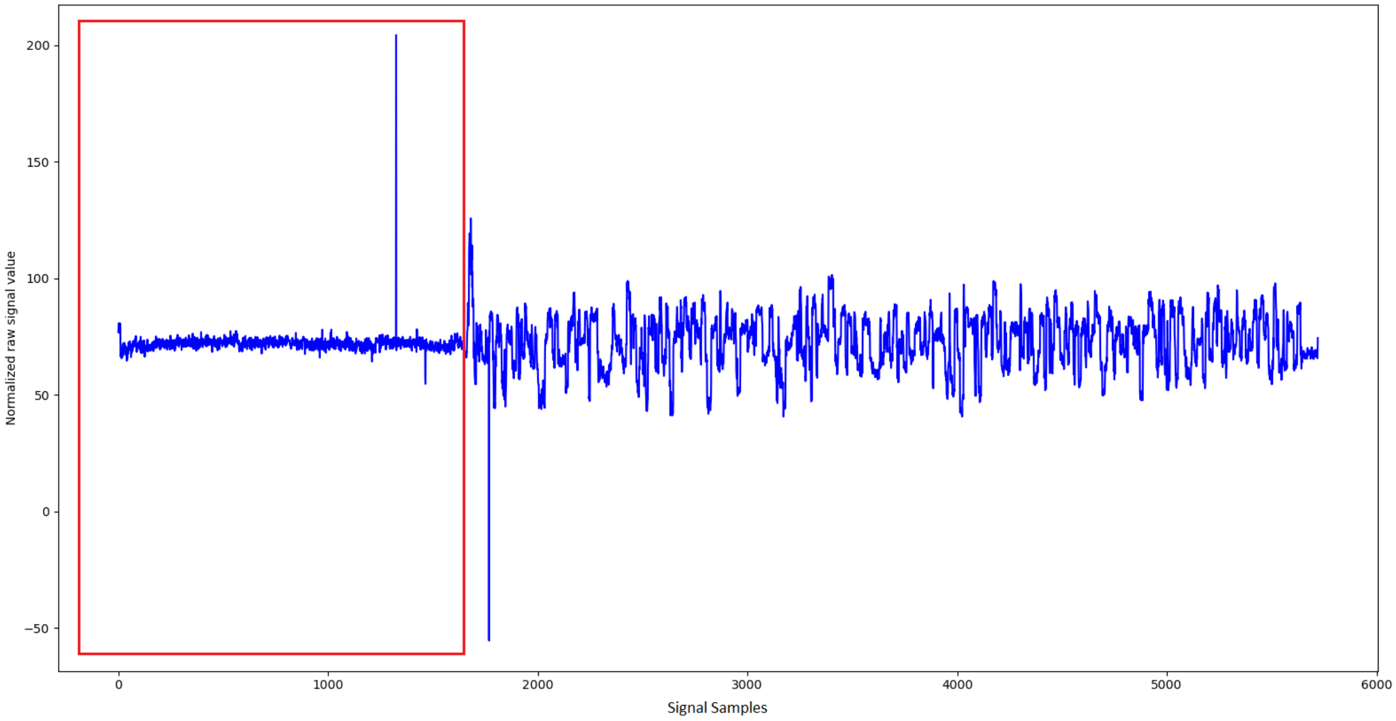

**Figure S1. Adapter segment in the first 1000-1500 signal samples in reads**

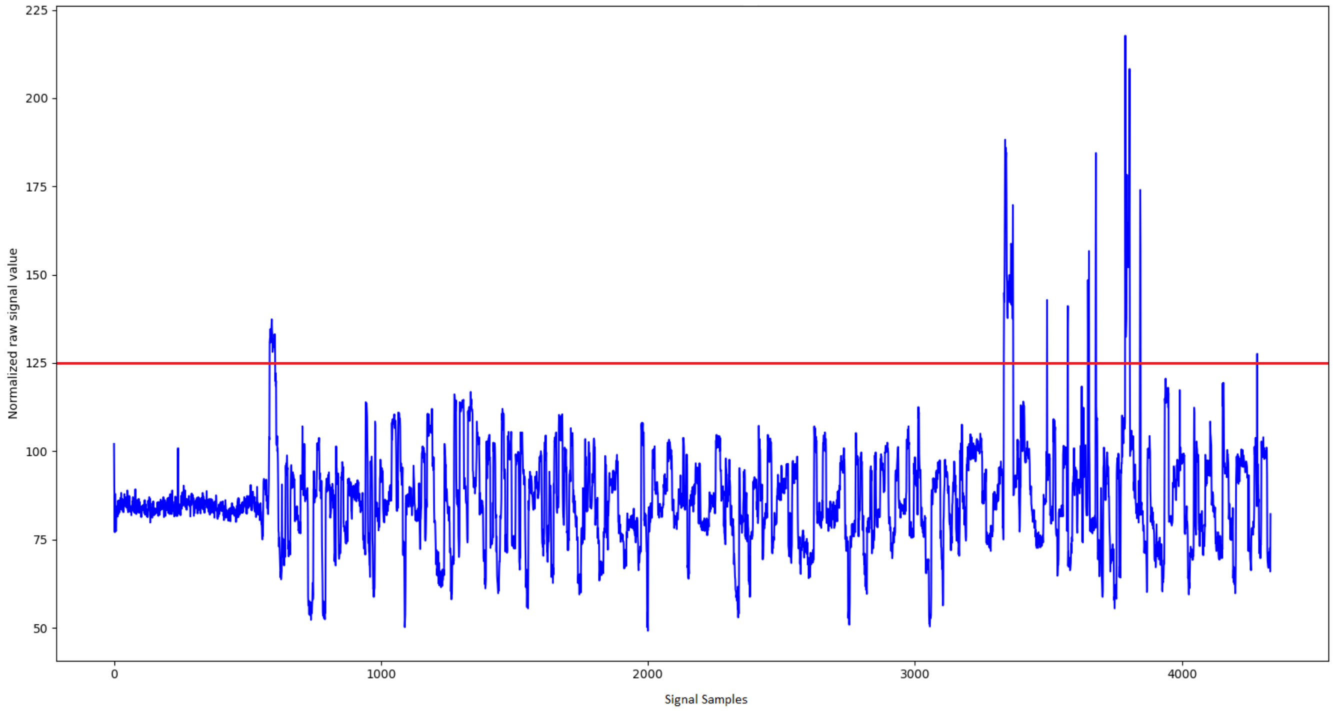

**Figure S2. Outliers in sequence signal**

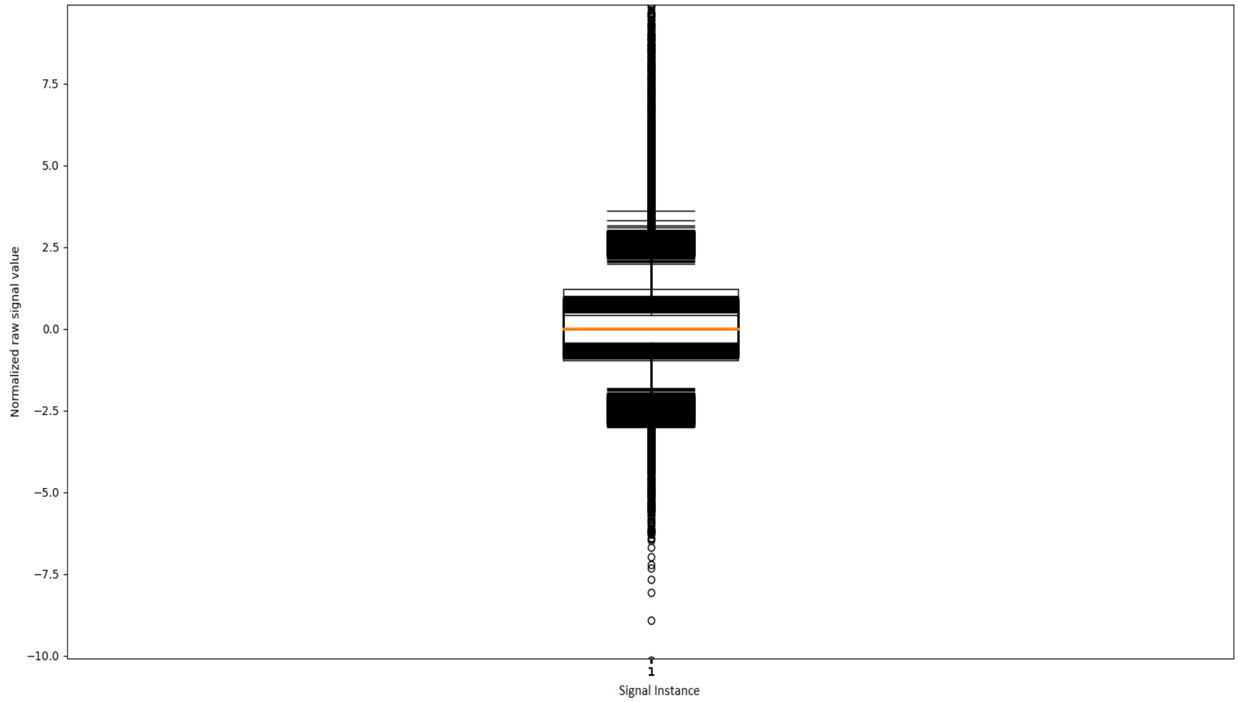

**Figure S3. Box plot of normalized raw signals for empirically deriving the optimal MAD threshold values**

**Supplementary Note 1 - Instructions to run the tools**

### Prerequisites

1. Download and install SLOW5 Tools toolkit.

```
#install HDF5 and zlib development libraries
sudo apt-get install libhdf5-dev zlib1g-dev

VERSION=v0.6.0

wget
"https://github.com/hasindu2008/slow5tools/releases/download/$VERSION/slow5too
ls-$VERSION-release.tar.gz" && tar xvf slow5tools-$VERSION-release.tar.gz &&
cd slow5tools-$VERSION/

./configure

make
```

2. Download dataset and extract. Let's call this directory `<DATASET_DIR>`.

```
wget https://doi.org/10.5281/zenodo.7111366
tar xvf DeepSelectNet_curated_datasets.tar <DATASET_DIR>
```

### DeepSelectNet

1. Download DeepSelectNet from Github repository.

```
git clone git@github.com:AnjanaSenanayake/DeepSelectNet.git

cd DeepSelectNet
```

2. Set up the environment for DeepSelectNet by following the instructions here.

3. Preprocess datasets with DeepSelectNet preprocessor.

```
python scripts/preprocessor.py -pos_s5 <DATASET_DIR>/COVID/train-covid -neg_s5
<DATASET_DIR>/ZYMO/train-zymo -b 20000 -c 1500 -sco 4 -mad 5 -o train-dump
```

4. Train the model with preprocessed datasets.

```
python scripts/trainer.py -d train-dump -s 0.7 -k 5 -e 200 -o trainedModel
```

5. Testing the model with the best-trained model.

```
python scripts/inference.py -model trainedModel/<best_model> -s5
<DATASET_DIR>/COVID/test-covid -lb 1 -mad 5 -o predicts-covid.txt

python scripts/inference.py -model trainedModel/<best_model> -s5
<DATASET_DIR>/ZYMO/test-zymo -lb 1 -mad 5 -o predicts-zymo.txt
```

**SquiggleNet**

1. Download DeepSelectNet from Github repository.

```
git clone -b slow5-support https://github.com/AnjanaSenanayake/SquiggleNet.git

cd SquiggleNet
```

2. Set up the environment for SquiggleNet by installing the package requirements here.

3. Splitting datasets to preprocess with SquiggleNet.

```
slow5tools view <DATASET_DIR>/COVID/train-covid | grep -v '^[#@]' | awk
'{print $1}' > read-ids-covid.txt

slow5tools view <DATASET_DIR>/COVID/train-zymo.blow5 | grep -v '^[#@]' | awk
'{print $1}' > read-ids-zymo.txt

cat read-ids-covid.txt | head 14000 > train-covid-squigglenet.txt
cat read-ids-covid.txt | tail 6000 > val-covid-squigglenet.txt
cat read-ids-zymo.txt | head 14000 > train-zymo-squigglenet.txt
cat read-ids-zymo.txt | tail 6000 > val-zymo-squigglenet.txt

mkdir train-covid-zymo
cp <DATASET_DIR>/COVID/train-covid.blow5 train-covid-zymo
cp <DATASET_DIR>/ZYMO/train-zymo.blow5 train-covid-zymo
slow5tools merge train-covid-zymo -o train-covid-zymo.blow5
```

```
mkdir test-covid-zymo
cp <DATASET_DIR>/COVID/test-covid.blow5 test-covid-zymo
cp <DATASET_DIR>/ZYMO/test-zymo.blow5 test-covid-zymo
slow5tools merge test-covid-zymo -o test-covid-zymo.blow5
```

4. Preprocess datasets with SquiggleNet preprocessor.

```
python preprocess.py -b 14000 -ft slow5 -gp train-covid-squigglenet.txt -gn
train-zymo-squigglenet.txt -i train-covid-zymo.blow5 -o output

python preprocess.py -b 6000 -ft slow5 -gp val-covid-squigglenet.txt -gn
cal-zymo-squigglenet.txt -i train-covid-zymo.blow5 -o output
```

5. Training datasets with SquiggleNet.

```
python trainer.py -tt output/pos_14000.pt -nt output/neg_14000.pt -tv
output/pos_6000.pt -nv output/neg_6000.pt -o trainedModel.ckpt
```

6. Training datasets with SquiggleNet.

```
python inference.py -m trainedModel.ckpt -ft slow5 -b 1000 -i
test-covid-zymo.blow5 -o predictions
```

#### Baseline

```
cd DeepSelectNet/support

sh baseline.sh <DATASET_DIR>/COVID/test-covid.blow5
<DATASET_DIR>/ZYMO/test-zymo.blow5 <DATASET_DIR>/COVID/test-covid.fastq
<DATASET_DIR>/ZYMO/test-zymo.fastq  <DATASET_DIR>/COVID/covid-ref.fasta
<DATASET_DIR>/ZYMO/zymo-ref.fasta
```

#### Guppy_hac+Minimap2

```
cd DeepSelectNet/support

sh baseline.sh <DATASET_DIR>/COVID/test-covid.blow5
<DATASET_DIR>/ZYMO/test-zymo.blow5 <DATASET_DIR>/COVID/test-covid.fastq
<DATASET_DIR>/ZYMO/test-zymo.fastq  <DATASET_DIR>/COVID/covid-ref.fasta
<DATASET_DIR>/ZYMO/zymo-ref.fasta 300
```

### Guppy_fast+Minimap2

```
cd DeepSelectNet/support

sh readfish.sh <DATASET_DIR>/COVID/test-covid.fastq
<DATASET_DIR>/ZYMO/test-zymo.fastq <DATASET_DIR>/COVID/covid-ref.fasta
<DATASET_DIR>/ZYMO/zymo-ref.fasta
```
